# Multiple Channel Electrocardiogram QRS Detection by Temporal Pattern Search

**DOI:** 10.1101/2021.08.15.456413

**Authors:** Bruce Hopenfeld

## Abstract

A highly constrained temporal pattern search (“TEPS”) based multiple channel heartbeat detector is described. TEPS generates sequences of peaks and statistically scores them according to: 1) peak time coherence across channels; 2) peak prominence; 3) temporal regularity; and 4) number of skipped beats. TEPS was tested on 31 records of three channel capacitive electrode data from the UnoViS automobile database. TEPS showed a sensitivity (SE) of 91.3% and a false discovery rate (FDR) of 3.0% compared to an SE and FDR of 75.3% and 65.0% respectively for a conventional single channel detector (OSEA) applied separately to the three channels. The peak matching window was 30ms. The percentage of 5 second segments with average heart rates within 5 beats/minute of reference was also measured. In 6 of the 31 records, TEPS’ percentage was at least 30% greater than OSEA’s. TEPS was also applied to synthetic data comprising a known signal corrupted with calibrated amounts of noise. At a fixed SE of 85%, increasing the number of channels from one to two resulted in an improvement of approximately 5dB in noise resistance, while increasing the number of channels from two to four resulted in an improvement of approximately 3dB in noise resistance. The quantification of noise resistance as a function of the number of channels could help guide the development of wearable electrocardiogram monitors.

## 1. Introduction

Frequently encountered noise sources confound electrocardiogram heart beat detection methods that rely solely on size or shape criteria to distinguish QRS complexes from noise peaks.^8^ A single channel convolutional neural network (CNN) described by Cai and Hu has shown some promise in overcoming the limitations of conventional peak size/shape based QRS detectors by implicitly exploiting temporal patterns in single channel normal sinus rhythm cardiac signals.^3^ However, CNNs often don’t generalize well, and the Cai and Hu detector has relatively poor temporal resolution, with each output node corresponding to 16 ms.^3^

Antink^1^ and Ravichandran^10^ independently extended neural networks to QRS detection in 3 channel capacitive electrode recordings from the UnoViS automobile database^14^ (described further below). Antink’s CNN architecture achieved a sensitivity of 88.0% with a false discovery rate of 4.8% based on a 150ms peak matching window. Ravichandran^10^ described an encoder-decoder architecture that generates from the 3 channels a single denoised signal, which is then provided to a conventional QRS detector. Ravichandran performed a heart rate variability analysis on a test set that consisted of 7 of the 31 UnoViS automobile records and reported that the resulting heart rate variability metrics were similar to the ground truth metrics. Again, it is not clear how these multiple channel neural network approaches will generalize to other data sets.

Hopenfeld^6^ described a scheme that overcomes the size/shape limitation by exploiting the relative temporal regularity of the true peaks in sinus rhythm. In particular, a search through a set of peak times generates temporally regular sequences, which are scored according to temporal regularity likelihood. However, Hopenfeld did not describe the application of this method to real world noise signals. Further, this method was not applied to multiple-channel signals.

Conventional multiple-channel QRS detectors improve on single channel detectors by providing some degree of redundancy but will tend not to perform much better than single channel detectors when high noise corrupts all channels. The redundancy results from performing QRS detection separately in each channel, and creating a consistent final sequence from the resulting single channel sequences through a variety of techniques.^12,13^ However, if all of the channels suffer from high noise such that none of the individual sequences has high quality, the fusion of these sequences according to existing methods will tend not to improve the final sequence compared to the individual sequences.

Figure 1 illustrates of a scenario that would likely confound conventional multiple-channel QRS detectors. The left panels in Figure 1 show a 5-second-long segment, comprising two signals recorded from capacitive electrodes, from the UnoViS automobile database.^14^ In particular, there are many relatively large noise peaks that have similar shapes as true peaks (indicated by triangles) in the right panels, which show filtered and differenced versions of the raw signals.

**Figure 1.**
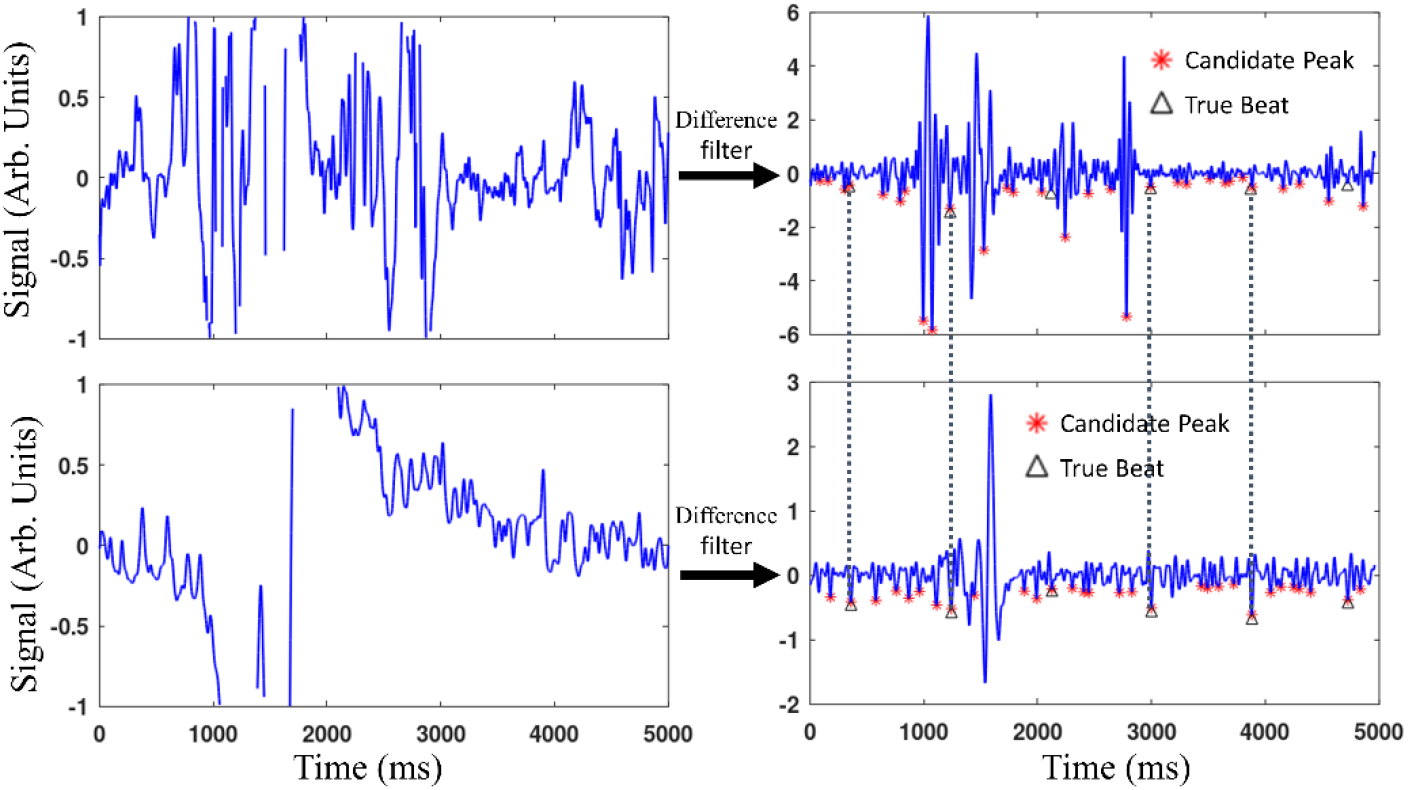
Raw (left panels) and filtered/differenced versions (right panels) of a five second segment from subject 6 of the UnoViS automobile database. The 5 second segment starts at 405 seconds from the beginning of the record. The top panel shows the average of the signals from channels 1 and 3, which were highly correlated; the bottom panel shows the signal from channel 2. The 30 most (negatively) prominent peaks are labelled by red asterisks and referred to as Candidate Peaks. The dashed black lines indicate Candidate Peaks in separate channels that temporally align, which suggests these peaks are more likely to be true beats.

This paper describes a methodology, “TEmporal Pattern Search” or “TEPS,” that can accurately detect the true peaks in this segment. The contributions of this paper include:

- The extension of sequence search and likelihood scoring to real world electrocardiogram signals
- The extension of sequence search and likelihood scoring to multiple channel signals
- The introduction of a likelihood score that depends on peak prominence
- The introduction of a likelihood score for multiple channel signals that depends on peak time coherence across channels
- Accurate detection of QRS complexes, with good temporal resolution (within 4ms of reference), in signals corrupted by high levels of noise characterized by many noise peaks that cannot be distinguished from QRS complexes based on peak size/shape criteria
- A demonstration of how detection sensitivity as a function of noise level depends on the number of channels

Octave (ver. 5.2.0) code for the below described algorithm is available for research purposes at https://github.com/Hopenfeld/Teps.

## 2. Algorithm

Figure 2 is a high-level block diagram of TEPS, which processes data in non-overlapping five second segments. For each segment, after pre-processing, it selects a certain number of peaks (“Candidate Peaks”) in each of multiple channels and creates a set of global peaks (“Global Peaks”) by grouping all the channels’ Candidate Peaks together, merging peaks that are temporally close across channels. It scores the likelihood of each Global Peak and selects the highest scoring ones (“Quality Peaks”) and then searches within these peaks for sinus rhythm sequences (“Parent Sequences”). Next, it fills in the gaps in the Parent Sequences with the remainder of the (lower quality) Global Peaks, thereby generating a set of Offspring Sequences, which are scored for quality. The final score of each Offspring Sequence is a weighted sum of its raw score (“SC”) and the raw scores of temporally matching sequences in the prior and next segment. The Offspring Sequence with the highest final score is selected.

**Figure 2.**
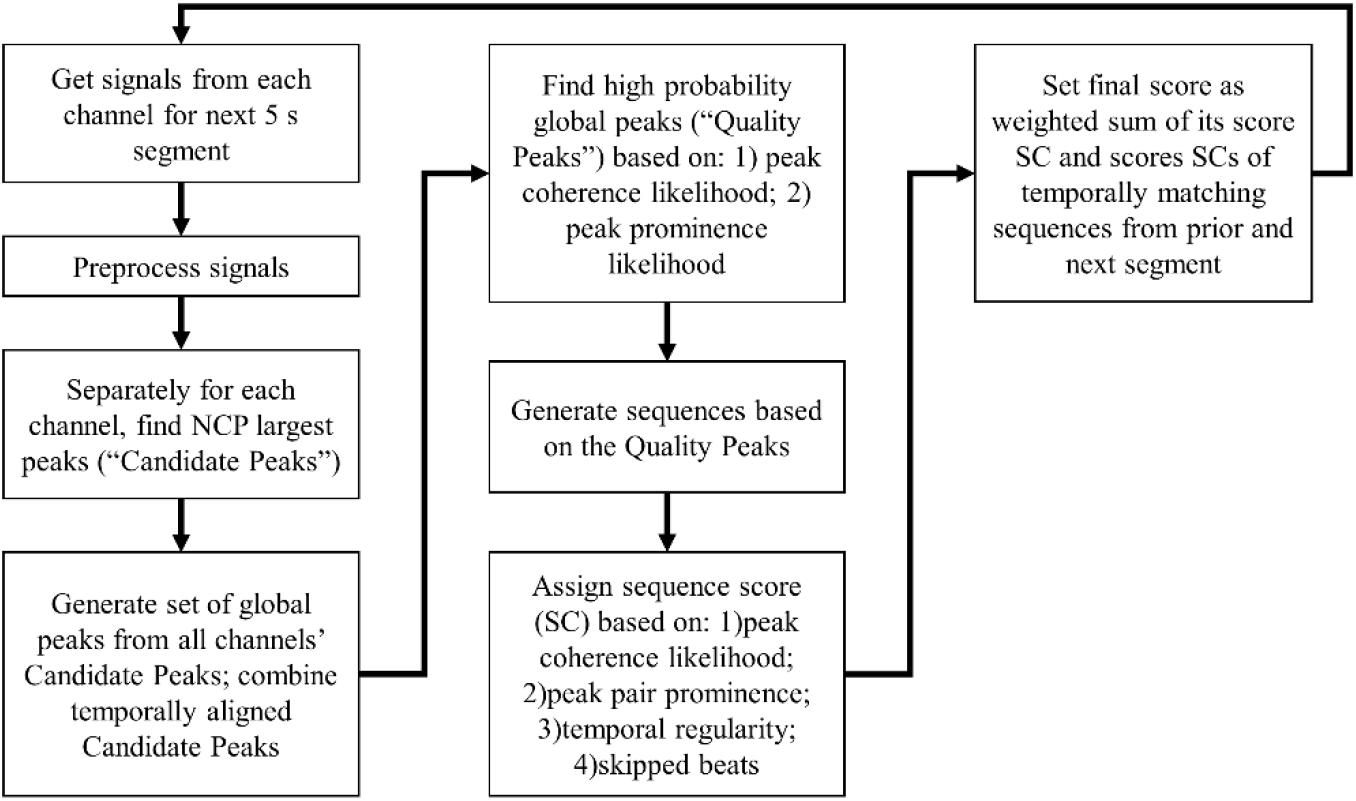
High level TEPS flowchart.

As described above, an assessment of likelihood is involved in both the selection of the Quality Peaks and the raw scoring of Offspring Sequences. The Quality Peak selection likelihood comprises two factors: 1) peak temporal coherence across channels; 2) peak prominence. The peak timing coherence score component is a Bayesian measure based on the tendency of true heart beats to occur close in time across channels whereas uncorrelated noise peaks will exhibit less coherence across channels. Figure 1 shows an example of this phenomenon. The peak prominence score component is based on the tendency of true heart beats to have greater magnitude than nearby peaks when compared to noise peaks.

Both factors also play a role in the scoring of each Offspring Sequence, which are further scored according to: 3) temporal regularity; and 4) the number of skipped beats. The temporal regularity score component is a measure that quantifies the change in between peak time intervals over a sequence. (The time between a sequence’s consecutive peaks will be referred to as an “RR interval.”)

Equations 1–8 below formalize the above mentioned four factors and set forth some of the statistically based motivation for aspects of the algorithm. However, as will be described, relevant probability distributions were generated based on limited data, and the search aspects of the algorithm complicate typical assumptions of independent events (in this case peaks). Although the generated probability distributions appeared to produce reasonable results for relatively high probability values, low probability values tended not to be reliable. For this reason, probability estimates are often summed, not multiplied together as would be mathematically appropriate given complete probability distributions. (For the purposes of obtaining a maximum probability solution, a multiplication rule corresponds to a logarithm summing rule, which penalizes low probability events compared to a regular summing rule.)

Returning to the selection of Quality Peaks from the set of Candidate Peaks, the likelihood score SCP for peak X is:

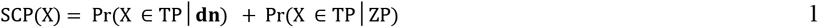

where TP indicates a true heart beat, Pr(X ∈ TP |**dn**) is the Bayesian peak timing coherence probability based on a set of peak time differences **dn**, and Pr(X ∈ TP |ZP) is the Bayesian peak prominence probability based on a peak prominence measure ZP. For each Offspring Sequence, the raw score SC is set equal to:

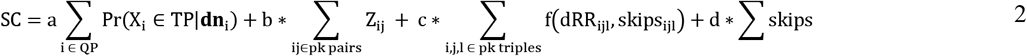

where Z is the prominence ratio for each peak pair in a sequence, dRR_ijl_ is the change in RR intervals between 3 consecutive peaks with corresponding skipped beats skips_ijl_, f() is a Gaussian with a standard deviation that depends on skips_ijl_. Thus, the peak coherence score of an Offspring Sequence, the first term on the right side of Equation 2, is simply the sum of the corresponding peak coherence scores of the Quality Peaks within the sequence. However, the sequence based peak prominence score, the second right hand side term, is based on a sequence level measure that is the sum peak pair prominence measures (Z_ij_). The relationship between the peak likelihood measure ZP and the peak pair ratios Z_ij_ will be described below.

A local peak will be referred to by “_x,α_” where the first subscript indicates channel number and the second subscript labels a Candidate Peak within a channel. A Global Peak will be referred by X or X_i_. Global Peaks are the union of the channels’ Candidate Peaks, with peak times set equal to the mean of the closely aligned local peak times:

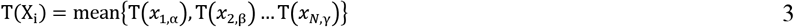

where T(x) is the peak time of peak x, and all of the local peak times T(x_a,α_) are separated by less than a time limit (e.g. 24 ms or 6 samples at a 256Hz sampling rate). If there are no local peaks in other channels within the time limit for a particular local peak, the global peak time is simply the local peak time.

### 2.1. Peak Coherence

Heart beats will occur close in time across channels whereas noise peaks (in uncorrelated channels) will tend to temporally align across channels only by chance. A Bayesian analysis of peak time coherence can therefore help to distinguish heart beats from noise. In the case of completely uncorrelated signals, the posterior probability of a Global Peak X corresponding to local peaks that are closest in time in respective channels occurring with time differences **dn** being heart beat peaks (TP=true peak) is approximately:

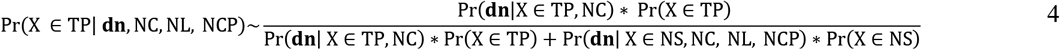

where NS denotes a noise peak, NC is the number of channels, NL is noise level, and NCP is the number of candidate peaks in each channel. The probability also depends on the number of samples in the search window, but since this quantity is held constant (5 seconds = 1280 samples), it is excluded from Equation 4 for convenience. The *a priori* probabilities of a peak being a true beat or noise depends in turn on NCP, NL and the heart rate. The noise level NL is not known although it can be estimated. The heart rate is not known although there may be a good estimate of it from another source or a recent, high quality prior segment. However, due to the limited amount of available noise data and to reduce algorithm complexity, the dependence on NL and heart rate is ignored.

Instead, the probability of interest is taken as Pr(X ∈ TP|**dn**, NC, NCP), the distribution of which was estimated by simulations. Specifically, as will be further described in Section 2.6, calibrated amounts of noise from channel 1 of the 30 minute long electromyogram noise record from Physionet Noise Tress Test Database^3,7^ (‘NSTDB’) was added to clean electrocardiogram signals, with known peak times, divided into 2, 3 and 4 channels. The resulting percentage of true peaks as a function of **dn**, NC and NCP was then tabulated.

### 2.2. Peak Pair Prominence

Compared to noise peaks, pairs of heart beats are more likely to be separated from one another by “smaller” peaks. Therefore, a Bayesian analysis of peak pair prominence can help to distinguish heart beats from noise. Figure 3 shows an example of the peak prominence of true beat pairs relative to noise pairs. The top and bottom panels of the figure show simultaneous differenced signals corresponding to first and second channels. True beats are designated by triangles and numbered by their order of occurrence. In channel 1, true beats 2 and 3 are more prominent than the intervening peaks. Likewise with true beats 6 and 7 in channel 2. Noise peaks separated by physiologically possible RR interval limits (e.g. between 300ms and 2000ms) tend to occur within bursts of noise and therefore tend to lack prominence relative to intervening noise peaks.

**Figure 3.**
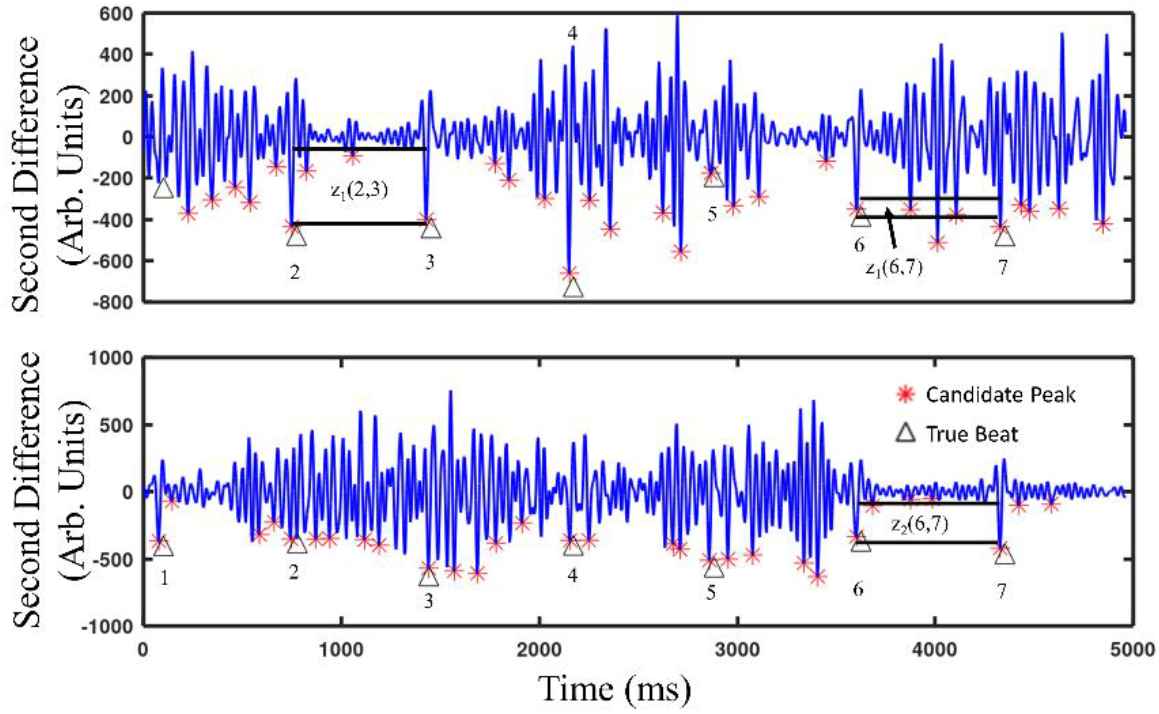
Examples of the peak pair prominence ratio Z. Channels 1 and 2 from the synthetic data are shown in the top and bottom panels respectively. The 7 peaks in a particular sequence are labelled 1 through 7. (Peak 1 does not exist in Channel 1 so is not labelled there.) The z ratio for a local peak pair is the ratio of the amplitudes of the lower and upper black bars respectively. Although only three z ratios are shown, the z ratio is calculated for every peak pair in the sequence for a total of 11 z ratios. As an example, the Z ratio for global peaks 6 and 7, Z(6,7), is z_1_(6,7)+z_2_(6,7).

The posterior probability of peak X characterized by a peak prominence measure ZP is approximately:

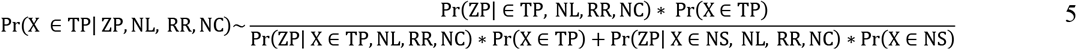

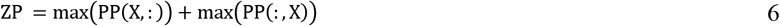

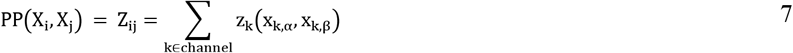

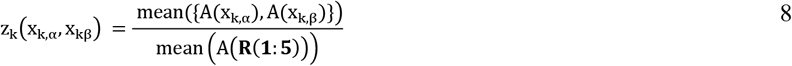

where x_k,α_ and x_k,β_ are the local peaks that correspond to global peaks X_i_ and X_j_ respectively, **R** is the set of the five largest peaks (not necessarily Candidate Peaks) in between x_k,α_ and x_k,β_; A() maps peak labels to peak amplitude (of the differenced signal); and sort indicates sorting in descending order. Again, to estimate the posterior probability, in this case Pr(X ∈ TP |ZP, NC, NCP), calibrated amounts of noise from channel 1 of the electromyogram noise record from the NSTDB was added to a clean signal.

The likelihoods are conditioned on the RR interval because the greater the time between peaks (RR interval), the more likely that larger noise peaks will occur between X_i_ and X_j_, so that Z will tend to be smaller. However, the conditioning on RR is not considered because of the relatively small size of the simulation data set. Instead, to take account of this RR effect, as will be described below, the probability of low average RR sequences is reduced according to a simple function.

The prominence based peak likelihood (Equation 5) affects selection of Quality Peaks according to Equation 1. In contrast, the sequence level summed Z scores, which is not a probability measure, factors directly into a sequence’s raw score according to the second term on the right-hand side of Equation 2. A sequence level Bayesian analysis of peak prominence is not performed due to the relatively small amount of training noise, which is nonetheless sufficient to produce reasonably smooth probability distributions at the peak level.

Summing Z scores across peak pairs will tend to benefit sequences with many peaks, which correspond to high heart rate sequences. Instead of normalizing summed Z scores based on the number of peaks, sequences with relatively low RR intervals are penalized by linearly decreasing Pr(X ∈ TP |ZP) with decreasing RR interval after it drops below a specified value (550ms).

The noise level NL, which is required for the estimate of Pr(X ∈ TP |ZP, NC, NCP), is not known *a priori*. For each segment in each channel, NL is estimated by analysing the distribution of peak prominences. The details are omitted because the algorithm’s results did not seem to be greatly affected by the accuracy of the noise estimate.

### 2.3. Temporal Regularity and Skipped Beats

The regularity of sinus rhythm can help distinguish true heart beat sequences from sequences of noise.^6^ The probability of finding a temporally regular noise sequence increases with the number of candidate peaks NCP and further depends on the characteristics of the noise: more uniformly distributed heart beat mimicking noise peaks will tend to result in sinus rhythm like sequences than more sporadically distributed noise. The *a posteriori* probability based on temporal regularity is thus a function of the noise type, NCP, and the *a priori* temporal regularity of sinus rhythm, which generally depends on RR interval.

Instead of implementing a Bayesian temporal regularity scheme, a temporal regularity measure TR, the third term on the right-hand side of Equation 2, forms a direct component of a sequence’s raw score according to Equation 2. In practice, it is not clear that a Bayesian measure would greatly improve the algorithm since the TR score, along with the skipped beats, tended to serve as a tie breaker amongst sequences whose primary quality was dictated by peak coherence probability and peak prominence.

To account for the greater temporal variability of low heart rate (high average RR interval) sequences,^11^ the TR scores for these sequences are increased according to a scaling factor that increases with RR interval above a certain threshold (e.g. 850 ms).

The Gaussian f(dRR_ijl_, skips_ijl_) for RR interval changes (=T(x_3_)−2*T(x_2_)+T(x_1_) for consecutive sequence peaks x_1_, x_2_, and x_3_) was generated^6^ from the Physionet Normal Sinus Rhythm RR Interval Database^3^ (‘NSRDB’), which consists of 24 h recordings of 54. The distributions for various numbers of skipped beats were generated by randomly eliminating peaks from the NSRdb records.

### 2.4 Channel Correlation

The correlation coefficient is computed for each pair of preprocessed signals. If the coefficient exceeds a threshold for a pair of signals, they are combined into one signal (and NC reduced by 1). Otherwise, the correlation coefficient linearly reduces the peak coherence probabilities Pr(X ∈ TP |ZP, NC, NCP) to reflect the relative lack of independence between channels.

### 2.5 Processing Details

Raw signals are preprocessed by low pass filtering with a 5^th^ order Butterworth filter with a cutoff frequency of 45Hz. The filtered signals are downsampled to 256Hz, and the resulting signal x() is differenced according to: y(i) = x(i-12)−2*x(i-6)+x(i). All computations were performed on a 2017 Hewlett Packard Laptop with an Intel Core i3-8130U CPU, base frequency 2.20GHz, with 8GB of RAM. The number of candidate peaks (NCP) is set equal to 25, 30, 35 and 40 for 1 through 4 channels, respectively. Candidate Peaks must meet broad minimum and maximum peak width criteria.

### 2.6 Databases; Testing and Parameter Fitting

The algorithm was tested on the UnoViS automobile dataset^14^ created by researchers at Medical Information Technology (MedIT), Helmholtz-Institute for Biomedical Engineering at RWTH Aachen University. This dataset consists of 3 channel ECG signals recorded from capacative electrodes attached to a car seat, along with a reference channel recorded from contact electrodes. The total duration of all records is over 13 hours.

To assess the relationship between detection performance, noise level, and number of channels, a synthetic data set was created. Specifically, channel 2 of the NSTDB electromyogram noise record was added to 1 to 4 copies of a synthesized clean signal whose RR intervals matched those of the first 30 minutes of record 1 of the NSRDB. (The NSTDB also contains a motion artifact noise record but this data was not considered suitable for the reasons described in Appendix 1.) These RR intervals range from approximately 450 ms to 735 ms. Because each segment of the noise data was added to a corresponding synthetic signal segment in a single channel, the durations of the 1-4 channel recordings were approximately 30, 15, 10 and 7.5 minutes respectively. Figure 4 shows examples of various noise levels. The noise level was determined by applying the Physionet nst script methodology (https://physionet.org/physiotools/wag/nst-1.htm) to a signal’s second difference (rather than the raw signal).

**Figure 4.**
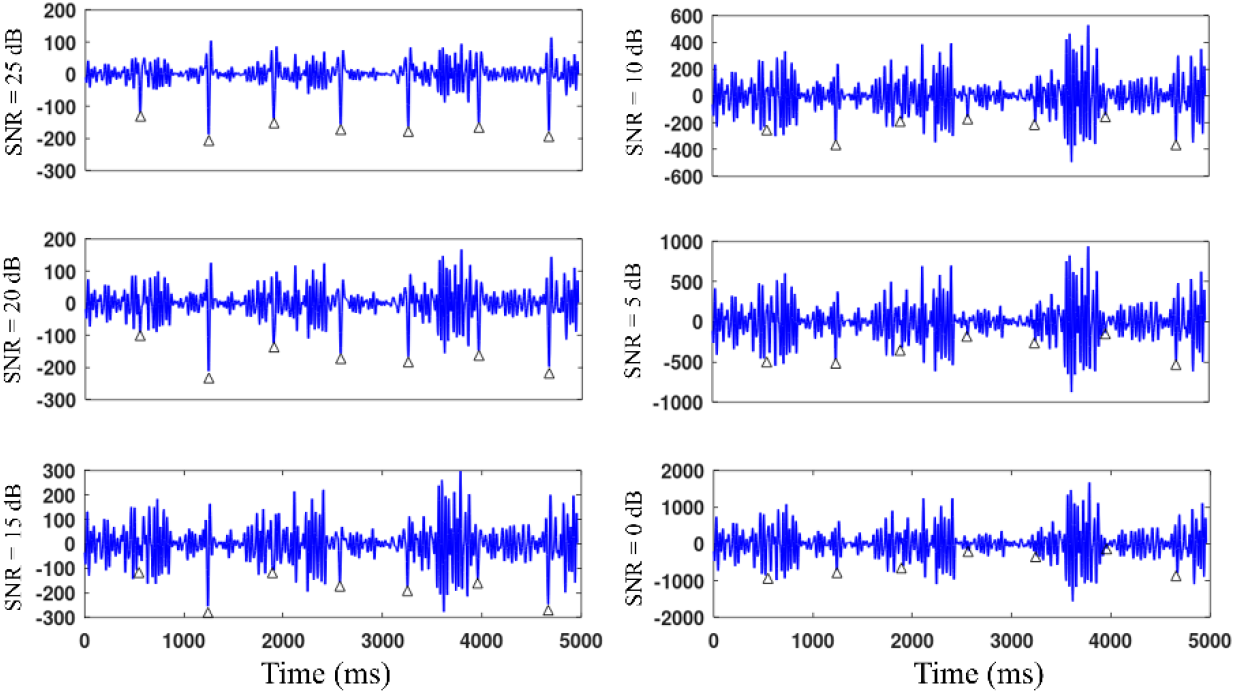
Examples of SNR values for the synthetic data. Triangles indicate true beats. Each panel is based on the same underlying clean segment.

The MedIT database has reference beat annotations that were generated by applying the open-source ECG analysis algorithm^5^ (OSEA) to the reference channel and the capacitive electrode channels. For the reference channel, a measure of the average RR interval over a 5 second segment was obtained from the annotations by (a) computing the interval between successive peaks, eliminating the intervals less than 500ms and greater than 1100ms, and taking the median of the resulting intervals if there were at least four of them; if so, the intervals were selected if (b) their standard deviation was less than 170ms. If these conditions were not satisfied, simple search of the peaks was carried out to find a regular sequence. If such could be found, then its mean RR interval was computed. (The data was exhaustively inspected to ensure that it was unlikely that any RR intervals were actually less than 500ms. TEPS is not limited to this RR interval range.)

To determine the difference between the OSEA heart rate in three channels and the reference heart rate, the channel that produced the closest match was taken. This likely at least somewhat overstates the match between the OSEA capacitive electrode annotations and the reference.

For all data, a peak was considered a match if it was within 15ms of the reference beat time.

The coefficients b,c, and d in Equation 2 were determined by performing a least squares fit of the data from record number 6 of the UnoViS automobile database. This record was chosen because it has a reliable reference signal and a good distribution of noise in the capacitive electrode channels. However, due to the frequent correlation of two of the three channels, there were frequently effectively only two channels available, which was not sufficient to generate good statistics for peak coherence probability (Equation 4). The corresponding coefficient “a” in Equation 2 was chosen heuristcally with reference to approximately 5 (5 second long) segments from the synthetic test data.

## 3. Results

The left panel in Figure 5 shows the fraction of 5 second segments with average heart rates within 5 beats/minute of the reference as estimated by both TEPS and OSEA for each of the 31 subjects in the UnoViS automobile data. In 6 of the 31 records (excluding record 6, which was used to fit coefficients as previously described), TEPS’ percentage was at least 30% greater than OSEA’s. For all the records excluding the 6^th^, (Equation 2), Sensitivity (“SE”) and False Discovery Rate (“FDR”) were as follows:

| Test Metric | TEPS | OSEA |
| --- | --- | --- |
| Sensitivity | 91.0% | 75.3% |
| False Discovery Rate | 3.0% | 65.0% |

**Figure 5.**
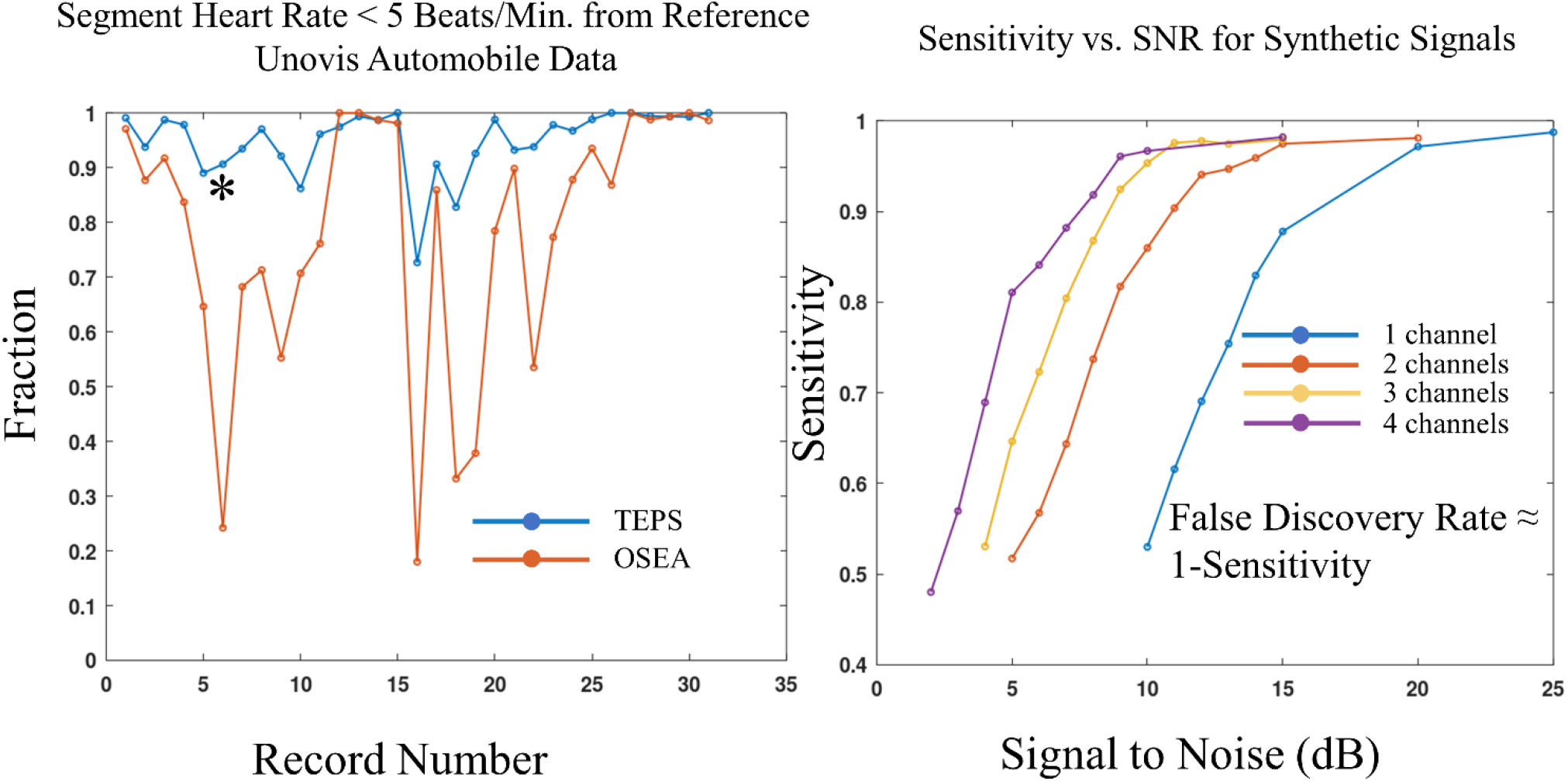
The left panel shows the fraction of average segment heart rates estimated by TEPS (blue) and OSEA (red) within 5 beats/minute of the reference value. The asterisk indicates that the sixth record was used to fit three TEPS coefficients and is therefore not strictly an appropriate test set record. The right panel shows sensitivity as a function of noise and number of channels for the synthetic data as processed by TEPS.

The mean time difference between TEPS and the reference for matching peaks was 1.26ms.

The right panel in Figure 5 shows the sensitivity of TEPS, as a function of SNR (Figure 4) for 1-4 channels of the synthetic noise data. In all cases, the FDR was approximately 1-SE, except for 1 channel at SNR levels below approximately 12dB, where the FDR was substantially larger. At a fixed SE of 85%, going from one to two channels resulted in an improvement of approximately 5dB in noise resistance, while going from two to four channels resulted in an improvement of approximately 3dB in noise resistance. The mean time difference between TEPS and the reference for matching peaks was 3-4 ms, depending on noise level.

## 4. Discussion

TEPS can accurately detect heart beats in conditions that would likely prove too noisy for conventional size/shape-based algorithms. The superior noise resistance of TEPS results from its exploitation of information that conventional algorithms either ignore or utilize in a superficial manner. Specifically, TEPS takes advantage of both heart rhythm and peak coherence across channels, which enables it to find patterns apart from peak size/shape.

Figure 6 is a stylized depiction of the benefit of the pattern finding approach. As shown, peaks are represented by circles whose diameters reflect peak size. The horizontal axis represents time while the vertical axis represents 3 different channels. Arrows point to the columns of true beats, whose regular pattern distinguishes them from noise peaks, despite the large overlap in size between the two types of peaks. Of course, if a size pattern exists, for example between the second and third true peaks in Channel 1, it can further help discriminate true beats from noise.

**Figure 6.**
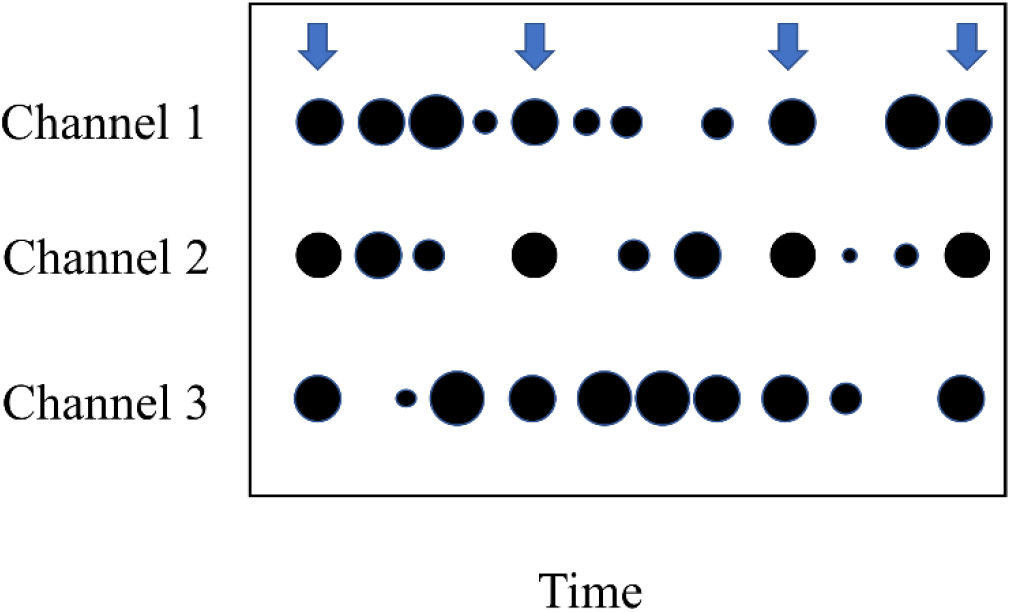
A stylized depiction of the pattern finding approach. Peaks are represented by circles whose diameters reflect peak size. The horizontal axis represents time while the vertical axis represents 3 different channels. Arrows point to the columns of true beats.

The temporal regularity of sinus rhythm and other heart rhythms implies a lower information entropy than irregular heart rhythms, which in turn means that regular rhythms can be detected in noisier channels than irregular rhythms. An optimal detector must take advantage of this regularity. However, TEPS can still detect irregular rhythms by performing another search after an initial search for sinus rhythm has failed to detect a high-quality sequence. The quality of the irregular rhythm sequence will depend on peak coherence likelihood and peak pair prominence, and possibility, characteristics of the rhythm to the extent it can be distinguished from noise.

Clinically valuable measures of heart rate variability can be extracted from somewhat noisy RR interval time series.^9^ With regard to TEPS, generation of a reliable RR time series would require excluding low quality segments, which in turn would benefit from an estimate of the accuracy of a detected sequence as a function of its final score. Assessing the relationship between detection accuracy and the final TEPS score is an area for future work.

It is somewhat difficult to compare the results of the present study with those of Antink^1^ (SE: 88.0%; FDR: 4.8%) because of the 120 ms difference in the peak time matching windows of the two studies. With regard to Ravichandran,^10^ it is simply not clear how well a denoising scheme would perform in very high noise conditions (e.g. 5dB in Figure 4) that TEPS can handle. Indeed, Ravichandran emphasized the applicability of the denoising system to cases where morphological information can be derived from the ECG,^10^ whereas TEPS is directed to high noise situations where only peak detection is feasible.

TEPS was tested on a very small data set, which limits the conclusions that can be drawn from this work. Yet, the minimal amount of parameter fitting that was done bodes well for its generalizability.

TEP was not tested on more than 2 channels in real world conditions since the UnoViS data often effectively comprised only 2 independent channels due to correlation. With only 2 independent channels, the peak prominence score (the second term in Equation 2) is frequently more important than the peak coherence score (the first term in Equation 2). The peak coherence score (and having more than 2 channels) becomes critical in extreme noise conditions, for example less than 5 dB (Figure 4), but TEPS was effectively tested against only the synthetic data in these conditions. For the synthetic data, the timing between peaks across channels between channels was a known quantity, 0 samples. Testing TEPS in real world, extreme noise conditions would require either calibration to ascertain this offset or additional processing to estimate it on the fly. These procedures may not produce a highly accurate estimate of this offset, which would degrade TEPS’ performance.

Yet another limitation of this study pertains to the comparison of TEPS with a single conventional algorithm, OSEA, which serially detects peaks. A search-based scheme that attempts to optimize the overall size and shape quality of detected beats^1^ may process this data more accurately than OSEA. Further, a more sophisticated multi-channel algorithm^13^ may have been able to reduce the FDR. However, for noisy segments such as those shown in Figure 1, any algorithm that ignores temporal information will likely fail.

## Acknowledgements

The author would like to thank James Hopenfeld and Dr. Hiroshi Ashikaga for comments on the manuscript.

## Appendix 1

TEPS was applied to both channels of the motion artifact noise record (‘em’) in the Physionet Noise Stress Test Database.^7^ A heart beat signal, with a normal rhythm in the range of 60 beats/minute, was detected throughout the entirety of both channels. A screen capture of a relatively clean 10 second segment shows a heart beat like signal in channel 1.

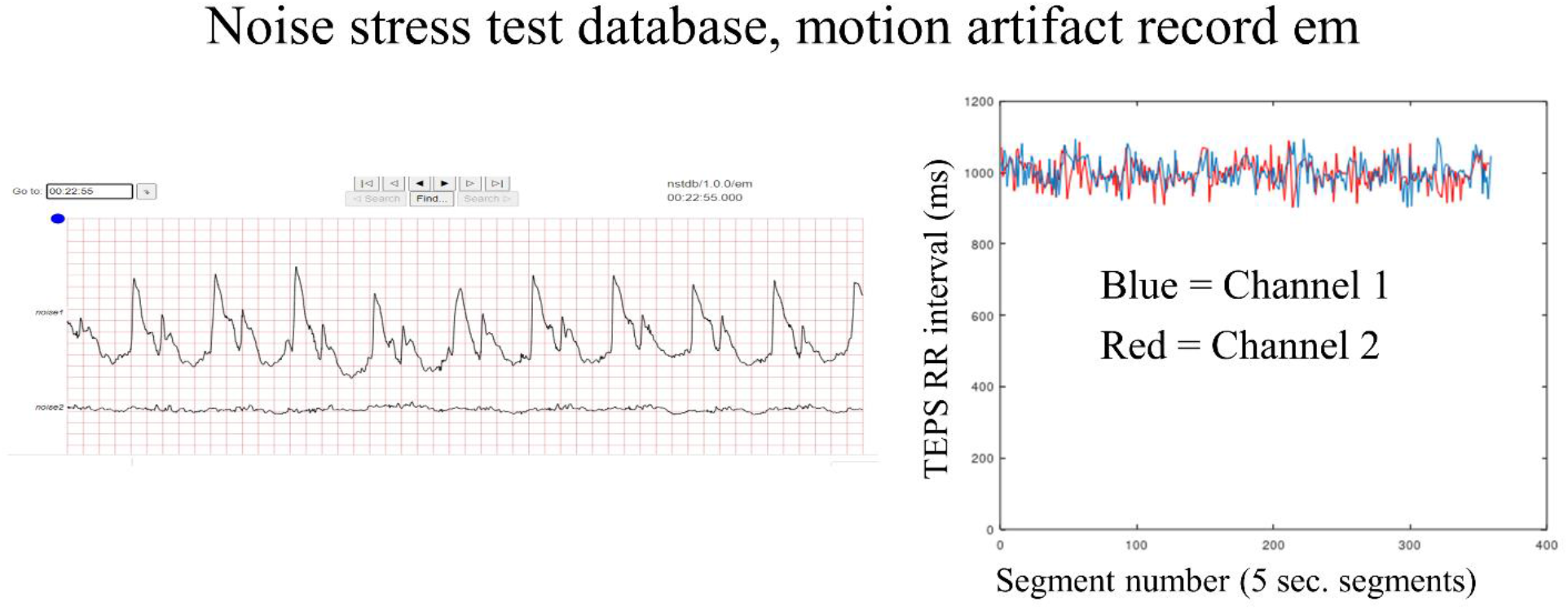

